# Voxel-wise Intermodal Coupling Analysis of Two or More Modalities using Local Covariance Decomposition

**DOI:** 10.1101/2022.02.19.481070

**Authors:** Fengling Hu, Sarah M. Weinstein, Erica B. Baller, Alessandra M. Valcarcel, Azeez Adebimpe, Armin Raznahan, David R. Roalf, Timothy E. Robert-Fitzgerald, Virgilio Gonzenbach, Ruben C. Gur, Raquel E. Gur, Simon Vandekar, John A. Detre, Kristin A. Linn, Aaron Alexander-Bloch, Theodore D. Satterthwaite, Russell T. Shinohara

## Abstract

When individual subjects are imaged with multiple modalities, biological information is present not only within each modality, but also between modalities – that is, in how modalities covary at the voxel level. Previous studies have shown that local covariance structures between modalities, or intermodal coupling (IMCo), can be summarized for two modalities, and that two-modality IMCo reveals otherwise undiscovered patterns in neurodevelopment and certain diseases. However, previous IMCo methods are based on the slopes of local weighted linear regression lines, which are inherently asymmetric and limited to the two-modality setting. Here, we present a generalization of IMCo estimation which uses local covariance decompositions to define a symmetric, voxel-wise coupling coefficient that is valid for two or more modalities. We use this method to study coupling between cerebral blood flow, amplitude of low frequency fluctuations, and local connectivity in 803 subjects ages 8 through 22. We demonstrate that coupling is spatially heterogeneous, varies with respect to age and sex in neurodevelopment, and reveals patterns that are not present in individual modalities. As availability of multi-modal data continues to increase, principal-component-based IMCo (pIMCo) offers a powerful approach for summarizing relationships between multiple aspects of brain structure and function. An R package for estimating pIMCo is available at: https://github.com/hufengling/pIMCo.

## 1 Introduction

There is increased availability of multi-modality neuroimaging data on individual subjects, with each modality containing unique information about brain structure or function. Such data allow us to explore patterns in individual modalities as well as patterns in the relationships between modalities, which we call intermodal coupling (IMCo), at global, regional, or local resolutions (Baller et al., 2021; Gu et al., 2021; Honey et al., 2009; Shokri-Kojori et al., 2019; Tak et al., 2015; Uddin, 2013; Valcarcel et al., 2018a, 2018b; Vandekar et al., 2016). The progress made by these IMCo studies have transformed our understanding of the brain, and it suggests that advancements in the methodology for studying IMCo have the potential to further enable such insights.

On the global scale, intermodal relationships have long been of interest. For example, structural connectivity (SC) and functional connectivity (FC) are strongly correlated in adults, but the relationship is less straightforward in children (Uddin, 2013). Gu et al. built on this global understanding of SC-FC coupling by exploring higher-resolution regional relationships – they found SC-FC coupling to be strongest in visual and subcortical areas with high SC (Gu et al., 2021).

In addition to structure-function relationships, regional relationships between metabolism and brain function have also been explored. In the study of energy utilization in the brain, Shokri-Kojori et al. showed that regional correspondence between cerebral glucose metabolism and fluctuations in blood oxygenation not only differed between brain networks in healthy patients but was also sensitive to differences between patients with acute or chronic alcohol use (Shokri-Kojori et al., 2019). Of note, these relationships were not identifiable by looking at individual modalities alone. Another regional study on metabolism-function coupling identified significant associations between cerebral blood flow (CBF) and strength of functional connectivity in default, frontoparietal, and primary sensory-motor networks. No significant association was found between CBF and functional connectivity strength in regions outside of these networks (Tak et al., 2015).

On the sub-regional local scale, studies from our group on coupling between cortical thickness and sulcal depth have suggested the cortical sheet is generally thinner in sulcal locations than in gyral locations and that this relationship was more spatially heterogeneous than previously described (Vandekar et al., 2016). A separate study exploring local IMCo between CBF and resting-state amplitude of low-frequency fluctuations (ALFF) showed that age-related declines in this measure of neurovascular coupling were most pronounced during mid-adolescence and were enriched in the dorsal attention network (Baller et al., 2021). There were also differences in CBF-ALFF coupling between males and females which were enriched in the frontoparietal network.

In these local IMCo studies, each vertex-wise coupling value was defined as the slope of the weighted linear regression (WLR) best-fit line for that local neighborhood between two modalities. However, because this method of calculating IMCo is based on regression slopes, it does not take into account vertex-level correlation and also suffers from inherent asymmetry, where coupling values depend on which modality is defined as the independent variable in the WLR. This asymmetry necessitates arbitrary, yet influential, choices when it comes to analysis and limits straightforward interpretation.

This measure for IMCo is also limited to only two modalities, so the study of coupling between more than two modalities using this method would require analysis of all pairwise couplings. As the number of total modalities increases, this approach can become challenging to interpret. Additionally, analysis of all pairwise couplings may not parsimoniously describe the overall degree of coupling across all modalities.

In response to these limitations, we propose a principal component analysis (PCA) based method for estimating IMCo that uses local covariance decomposition to define symmetric voxel-wise coupling values valid for two or more modalities. This method reduces complex local covariance structures into a single value, thus providing an easily interpretable value that characterizes the strength of coupling in settings with two modalities. It also allows for simplified study of more complex local covariance structures in settings with more than two modalities.

To demonstrate its sensitivity to biologically relevant patterns, we show that PCA-based IMCo (pIMCo) uncovers differences in three-modality coupling between CBF, ALFF, and regional homogeneity (ReHo) with respect to age and sex in youth. We chose these modalities because local cortical coupling between vascular organization and resting state fluctuations has been previously characterized. Additionally, while resting state fluctuations can be thought of as a more global property, regional homogeneity is more related to local connectivity. Thus, we were interested in understanding the coupling between cerebral blood flow and both long-range fluctuations and local connectivity.

## 2 Methods

### 2.1 Subjects

We included 803 subjects (340 males) from ages 8-22 (mean = 15.6; sd = 3.3) from the Philadelphia Neurodevelopmental Cohort (PNC) (Satterthwaite et al., 2014). Of the 1,601 PNC subjects who underwent neuroimaging, health screening as well as automated and manual image quality screening were performed. We excluded subjects in the following order: low T1-weighted MRI quality (n = 61), low resting-state fMRI (rfMRI) quality (n = 450), and low arterial spin labeling (ASL) quality (n = 54). Of the remaining subjects, we then excluded those meeting any of the following health exclusion criteria (n = 205): history of psychoactive medication, history of inpatient psychiatric hospitalization, or history of medical disorders that could impact brain function. Finally, ASL scans for which high-quality partial volume correction could not be performed were excluded (n = 28). This resulted in the final set of 803 subjects used for this study.

The Institutional Review Boards of the University of Pennsylvania and the Children’s Hospital of Pennsylvania approved all study procedures. All adult study subjects gave written informed consent; for subjects under the age of 18, parents or guardians provided written informed consent and subjects provided assent. Additional details of the PNC study have been previously described (Gur et al., 2020; Satterthwaite et al., 2014).

### 2.2 Image acquisition

All PNC imaging was acquired using a single 3T Siemens Tim Trio scanner with a 32-channel head coil. To minimize motion, subjects’ heads were stabilized using one foam pad over each ear and one foam pad over the top of the head. Image acquisition procedures have been previously described (Gur et al., 2020; Satterthwaite et al., 2014).

T1-weighted structural images were used for alignment of all scans into a common space. T1-weighted images were acquired using a 3D-encoded magnetization-prepared, rapid-acquisition gradient echo (MPRAGE) T1-weighted sequence with the following settings: *T*_*R*_ = 1810 ms; *T*_*E*_ = 3.51 ms; FoV = 180 × 240 mm; matrix size = 192 × 256; number of slices = 160; slice thickness = 1 mm; inter-slice gap = 0 mm; resolution = 0.9375 × 0.9375 × 1 mm. Cerebral blood flow (CBF) was estimated from a pseudo-continuous arterial spin labeling (pcASL) sequence with a spin-echo echoplanar readout and the following settings: *T*_*R*_ = 4000 ms; *T*_*E*_ = 15 ms; FoV = 220 × 220 mm; matrix size = 96 × 96; number of slices = 20; slice thickness = 5 mm; inter-slice gap = 1 mm; resolution = 2.3 × 2.3 × 6 mm; 80 volumes. Maps of amplitude of low frequency fluctuations (ALFF) and regional homogeneity (ReHo) were estimated from six minutes of task-free functional data from a blood-oxygen-level-dependent (BOLD) weighted 2D EPI sequence with the following settings: *T*_*R*_ = 3000 ms; *T*_*E*_ *=* 32 ms; FoV = 192 × 192 mm; matrix size = 64 × 64; number of slices = 46; slice thickness = 3 mm; inter-slice gap = 0 mm; resolution = 3 mm isotropic; 124 volumes. Subjects were instructed to stay awake, keep their eyes open, fixate on a displayed fixation cross, and remain still.

### 2.3 Image processing

Image processing of T1-weighted structural images, pcASL scans, and rfMRI scans have been previously described (Baller et al., 2021; Gur et al., 2020). They are summarized here in brief. T1-weighted structural images were processed using tools from Advanced Normalization Tools (ANTs) (Tustison et al., 2014). pcASL and rfMRI scans were processed using an eXtensible Connectivity Pipeline (XCP) which included tools from FSL and AFNI (Ciric et al., 2018; Cox, 1996; Jenkinson et al., 2012).

CBF was quantified from control-label pairs using the following equation:

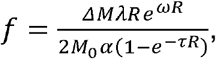

where f is CBF, *ΔM* is the difference in signal between control and label acquisitions, *R* is the longitudinal relaxation rate of blood, *τ* is the labeling time, *ω* is the post-labeling delay, *α* is the labeling efficiency, *λ* is the blood-tissue-water partition coefficient, and *M*_0_ is approximated by the control image intensity. We set *α* = 0.85, *λ* = 0.9*g*/*mL, τ* = 1.6*s, ω*= 1.2*s*. Partial volume correction was performed using Bayesian Inference for Arterial Spin Labeling MRI (BASIL) (Chappell et al., 2011; Chappell et al., 2009).

For rfMRI processing, the XCP pipeline included: 1) field inhomogeneity correction with FSL FUGUE, 2) removal of initial rfMRI volumes, 3) alignment of volumes within the time series to a selected reference volume using FSL MCFLIRT, 4) interpolation of intensity outliers with AFNI 3dDespike, and 5) demeaning and removal of linear or quadratic trends. Images were then denoised using a 36-parameter confound regression model that has been shown to minimize impact of motion artifact (Ciric et al., 2017). Finally, BOLD-weighted time series as well as artifactual model time series were filtered using a first-order Butterworth filter with a passband between 0.01 and 0.08 Hertz.

Voxel-wise ALFF was defined as the sum of frequency bins between 0.01 and 0.08 Hertz using a Fourier transform of the time-domain signal (Yang et al., 2007). Voxel-wise ReHo was defined as Kendall’s coefficient of concordance computed over the rfMRI time series in each voxel’s 26-voxel local neighborhood (Zang et al., 2004). Voxel-wise maps were smoothed with a 6mm full width at half maximum (FWHM) kernel to improve signal-to-noise ratio. CBF, ALFF, and ReHo images were co-registered to the the T1-weighted structural image using boundary-based registration and then normalized to a custom adolescent template using the top-performing SyN registration provided by ANTs (Avants et al., 2011; Ciric et al., 2021; Greve and Fischl, 2009). Finally, a gray matter mask was generated as the intersection between a gray matter mask from T1-weighted images with 90% coverage over all subjects and overall coverage masks from registered pcASL and rfMRI scans.

### 2.4 Methodology for estimating pIMCo

For each subject, we calculated voxel-wise coupling between CBF, ALFF, and ReHo images to produce one pIMCo image per subject. The full pIMCo estimation pipeline is summarized in Figure 1. First, we applied the gray matter mask to each of the three modalities. Then, within each masked modality, we globally scaled intensities to a mean of 0 and a variance of 1. This scaling is necessary because eigendecomposition is later performed on local covariance matrices; if modalities are defined on drastically different scales, decomposition outputs would reflect differences in scale between modalities rather than local covariance structures. Next, for each voxel, we extracted local neighborhoods from each of the three modalities and weighted voxels within these local neighborhoods proportional to a Gaussian kernel over their Euclidean distances from the central voxel – in our study, we used FWHM = 3, which corresponds to 7×7×7 voxel (14×14×14 mm) local neighborhoods and a standard deviation of 1.62 mm for the Gaussian kernel. Then, we calculated the 3×3 weighted covariance matrix between the neighborhoods, performed eigendecomposition on it, and extracted the proportion of variance explained by the first eigenvalue. Once all the voxel-wise proportional first eigenvalues were extracted, we scaled these values such that their theoretical range was [0, 1] and performed a logit transformation. While such a transformation makes the coupling value more challenging to interpret, it emphasizes extreme values of coupling and changes the domain of coupling values from 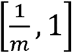 to (− ∞, ∞), where *m* is the total number of modalities, in order to improve expected behavior with post-hoc voxel-wise statistical analyses. This resulted in our voxel-level pIMCo image for that subject. For any particular voxel in that image, a large value suggests that the voxel’s local covariance matrix across modalities could be well-summarized in a single dimension while a small value suggests multiple dimensions would be necessary to characterize the covariance structure.

**Figure 1:**
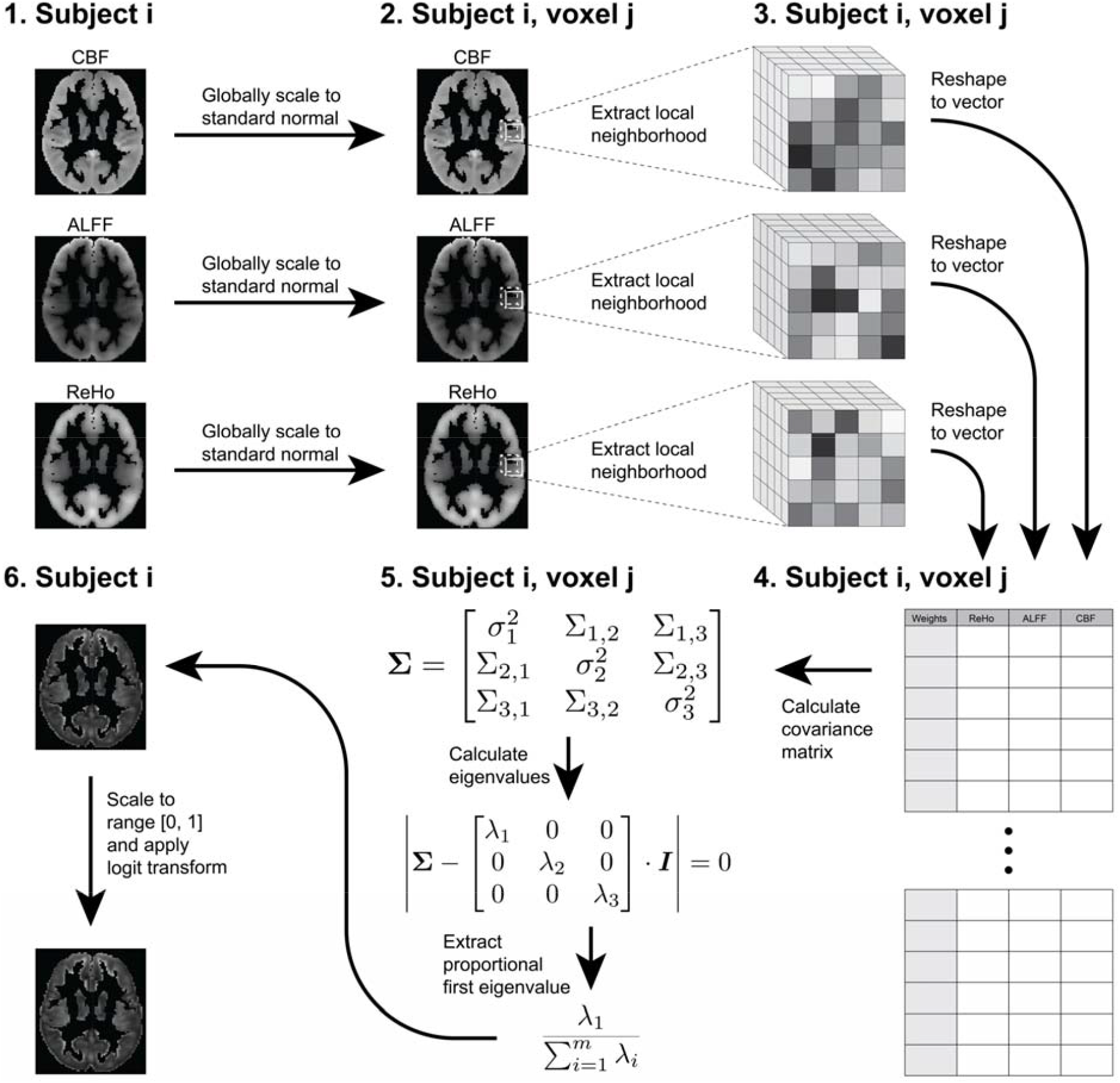
Step-by-step diagram of pIMCo estimation pipeline as described in the Methods. Coupling images are generated from intermodal images for each subject individually. pIMCo estimation is performed at each voxel location across subject-specific images.

For reference, in the three-modality setting, coupling values of -2, 0, and 2 correspond to the first eigenvector explaining 41%, 67%, and 92% of the total variance in that local neighborhood, respectively. In the two-modality setting, coupling values of - 2, 0, and 2 correspond to the first eigenvector explaining 56%, 75%, and 94% of the total variance in that local neighborhood, respectively.

### 2.5 Voxel-wise statistical analysis

We created descriptive coupling maps by taking the means and variances across all 803 subjects’ pIMCo values at each voxel location in volumetric space. We then projected these mean and variance images to the cortical surface using PySurfer for visualization of spatial heterogeneity and cortical patterns (Waskom et al., 2020).

To investigate the biological relevance of pIMCo, we used linear regression at each voxel to explore whether coupling was associated with age or sex. In all linear regressions, we controlled for in-scanner motion for both ASL and rfMRI scans. To account for multiple comparisons in these voxel-level tests, we controlled the false discovery rate at 5% (Benjamini and Hochberg, 1995). Then, we created binary thresholded masks indicating which voxels displayed a significant effect for each of age and sex. For this and following analyses, we performed identical modeling of each of the three modalities individually to explore whether age and sex effects were present and corresponded to the observed associations with pIMCo.

### 2.6 Spin testing

To visualize the extent of voxels where coupling was associated with age and sex, we counted the proportion of voxels with statistically significant age or sex effects in each of the Yeo 7 functional networks on the cortex as well as in subcortical regions in the Automated Anatomical Labeling (AAL) atlas (Thomas Yeo et al., 2011; Tzourio-Mazoyer et al., 2002).

Next, we tested whether the proportion of significant voxels in each functional network was enriched when compared to the proportion of significant voxels overall. Because there is an underlying spatial distribution of significant voxels, we used the spin test (Alexander-Bloch et al., 2018). Briefly, the spin test is a permutation-inspired testing procedure that rotates the FreeSurfer sphere randomly to create an underlying null distribution that preserves spatial patterns. The null hypothesis is that there is no spatial enrichment of significant p-values in the specified functional network compared to across the cortex overall. In our study, we estimated the null distribution over 2,000 permutations – for each permutation, we recorded the Jaccard similarity index between the thresholded p-value map and each of the Yeo 7 networks. Finally, for each network, we calculated the p-value as the proportion of null Jaccard similarity indices equal to or greater than the observed Jaccard similarity index.

### 2.7 Code and data availibility

An R package for calculating pIMCo images is available at: https://github.com/hufengling/pIMCo. All code for analysis is available at: https://github.com/hufengling/IMCo_analyses. The following software and packages were used for pre-processing, pIMCo calculation, or analysis – R Core Team (2021); Wickham (2019); Xie (2021a); Xie (2021b); Allaire et al. (2021); Xie (2016); Wickham et al. (2019); Muschelli (2021a); Avants (2020); Kandel et al. (2020); Müller (2020); Fischl (2012); Schäfer (2021); Jenkinson et al. (2012); Muschelli et al. (2015); Ren (2021); Muschelli (2021b); Yushkevich et al. (2006).

PNC data are publicly available in raw format at https://www.ncbi.nlm.nih.gov/projects/gap/cgi-bin/study.cgi?study_id=phs000607.v3.p2.

## 3 Results

### 3.1 Coupling varies across the cortical surface and is increased in discrete regions, bilaterally

We calculated voxel-wise mean and variance maps of coupling values to characterize spatial patterns in CBF-ALFF-ReHo coupling and visualized these on the cortical surface. Throughout the cortical surface, all voxels, on average, showed strong coupling, and voxels with stronger average coupling also tended to have higher variance between subjects (Pearson correlation = 0.69). The average voxel-wise mean coupling value was 0.99 (sd = 0.37; range = [0.27,3.30]). The average voxel-wise variance was 0.91 (sd = 0.20; range = [0.45,2.64]).

Visual comparison of voxel-wise descriptive maps with the Desikan-Killiany cortical atlas (Desikan et al., 2006) suggested that coupling is especially strong in the following regions, bilaterally: superior frontal gyrus, paracentral gyrus, caudal anterior cingulate, posterior cingulate, isthmus cingulate, pericalcarine, lateral occipital, and insula (Figure 2). Comparing to the Yeo 7 functional networks, areas of strong coupling are observed primarily in the frontoparietal (p = 0.0125) and default networks (p = 0.039; Figure 3).

**Figure 2:**
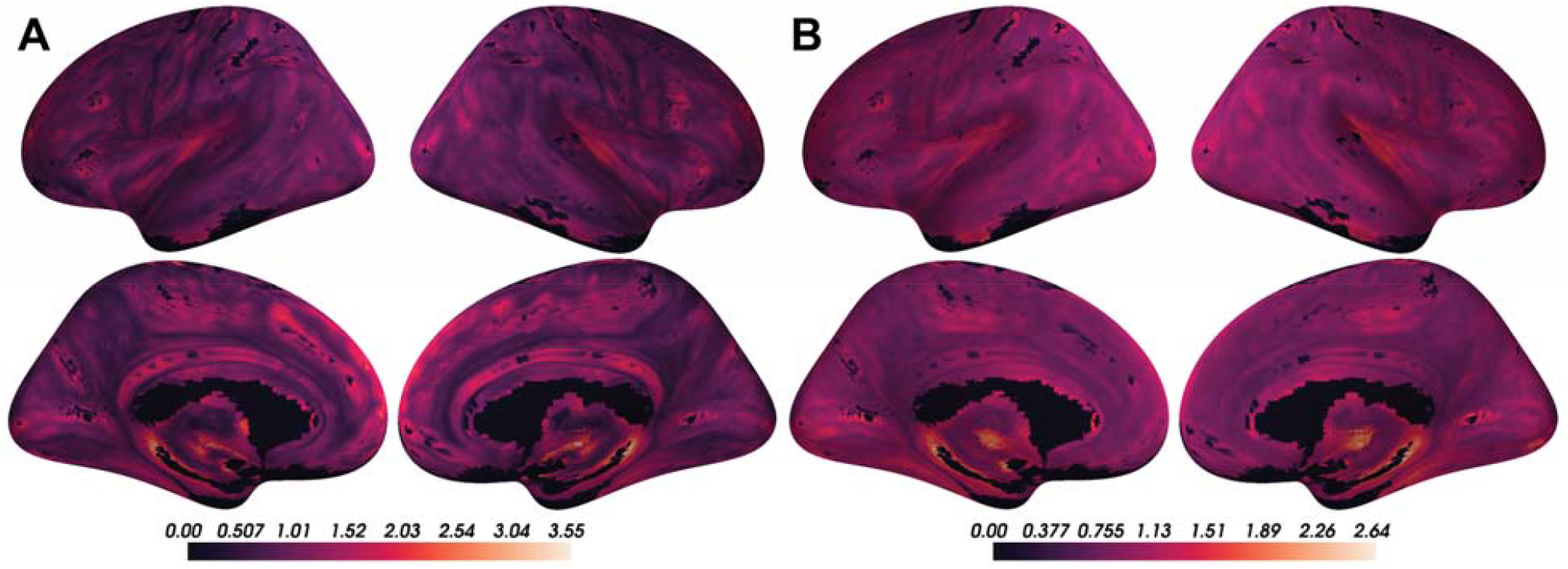
Coupling values are spatially heterogeneous across the cortical surface. A) Voxel-wise means across subjects of cortical coupling values between CBF, ALFF, and ReHo. Larger values indicate stronger coupling. B) Voxel-wise variances across subjects of cortical coupling values between CBF, ALFF, and ReHo.

**Figure 3:**
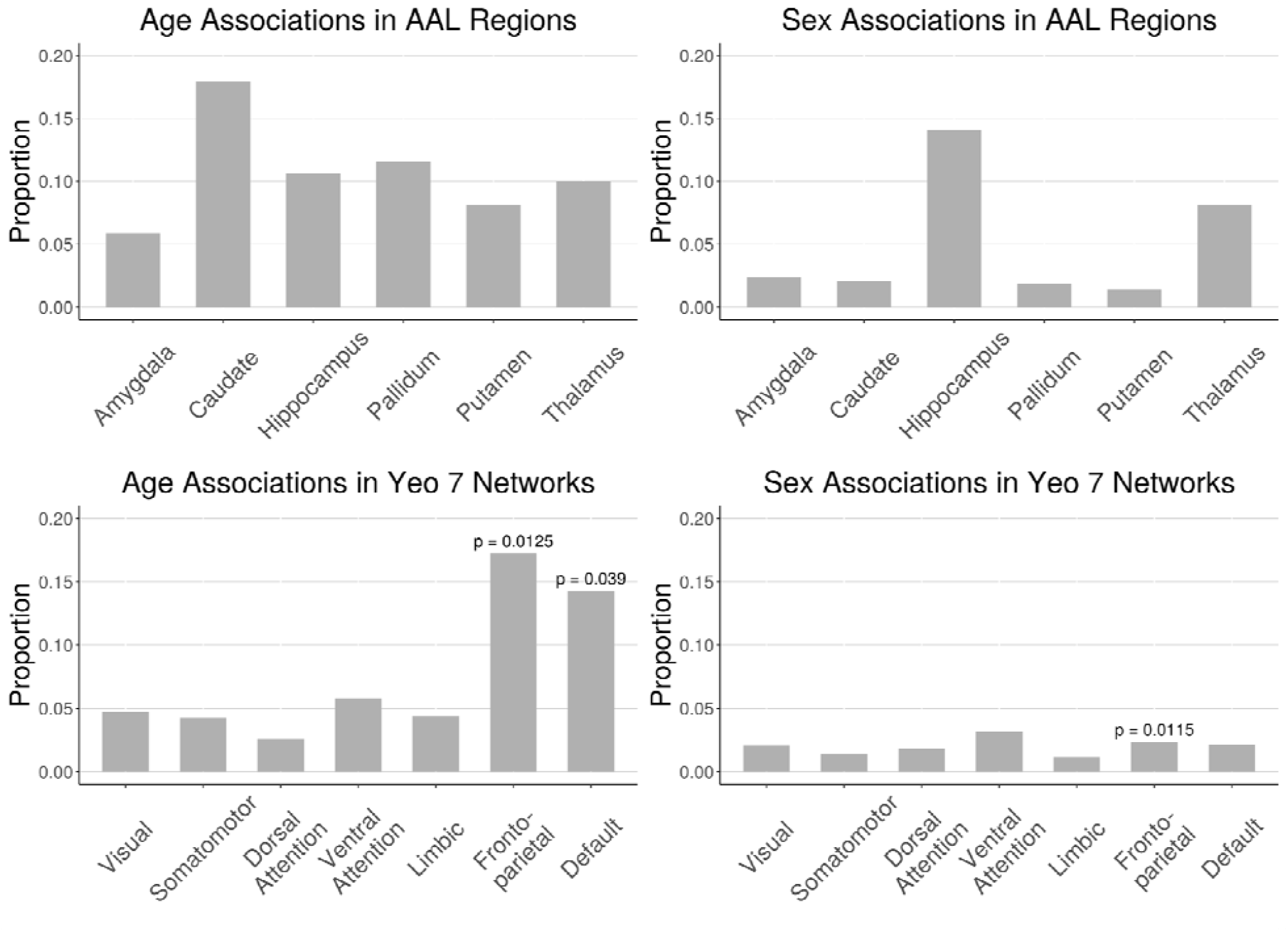
Proportion of voxels in AAL subcortical regions and Yeo 7 cortical networks that showed significant coupling-age and coupling-sex associations when in-scanner motion was included as a covariate (FDR corrected p < 0.05). Spin test was performed for all Yeo 7 networks; significant p-values are reported (p < 0.05).

These voxel-wise descriptive maps of coupling showed unique information when compared to the descriptive maps of each of the individual modalities (Supplementary Figure S1).

### 3.2 CBF-ALFF-ReHo coupling evolves with age throughout gray matter structures

Linear associations between strength of coupling and age were present in subcortical structures and cortical networks (Figure 3; corrected p < 0.05). Figure 4 shows an example of such an association between coupling and age as well as individual modalities and age at one voxel in the default network. In subcortical structures, age-related changes in CBF-ALFF-ReHo coupling occurred primarily in the caudate and pallidum, though such changes were also common in the hippocampus, putamen, and thalamus.

**Figure 4:**
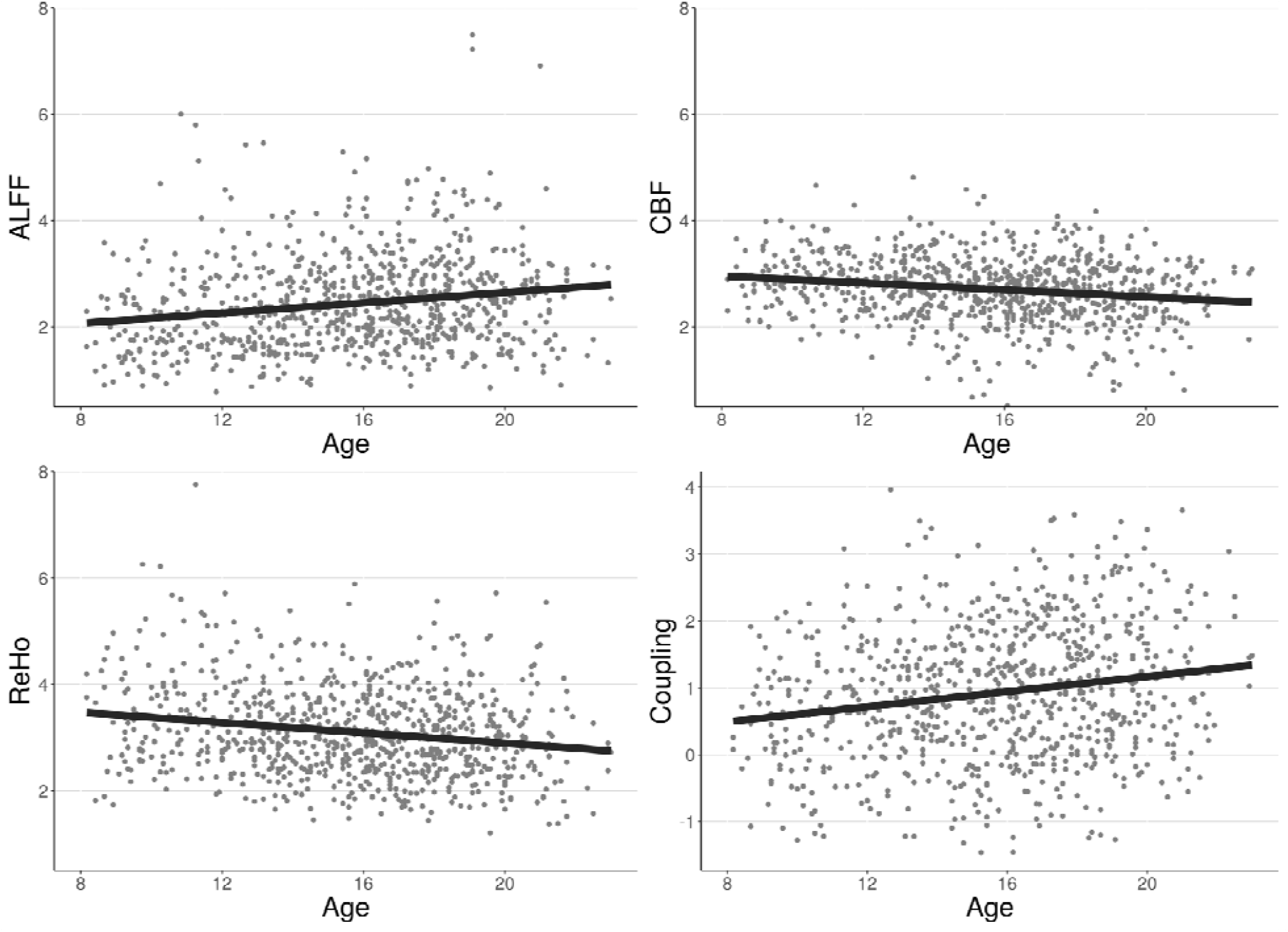
Example of associations between individual modalities and age as well as associations between coupling and age at a single voxel in the default network. Each point represents the value at that voxel for one subject. Best-fit lines from univariate linear regression are shown.

In cortical networks, coupling and age associations were rare in all networks except the frontoparietal and default networks (Figure 5). These two networks were also the networks in which the average strength of coupling across subjects appeared to be highest (Figure 2). Spin testing between functional networks and age-related changes in coupling showed enrichment of coupling and age associations in the frontoparietal (p = 0.013) and default networks (p = 0.039).

**Figure 5:**
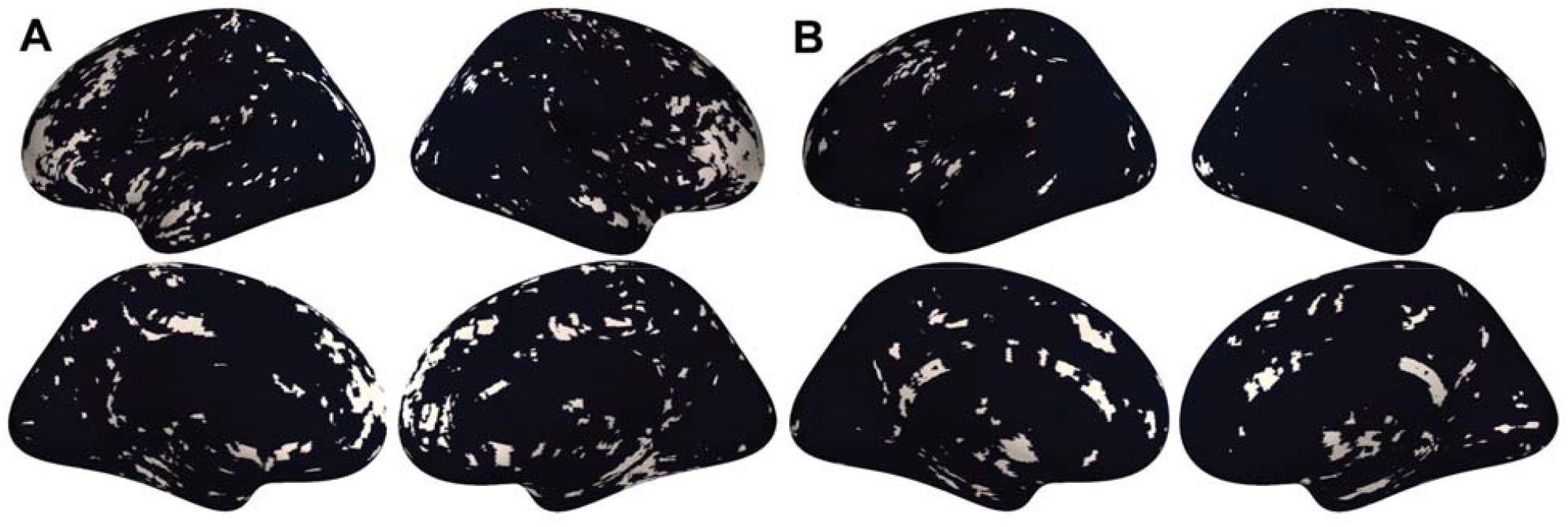
A) Thresholded maps of voxels with significant coupling associations with age after FDR correction at 0.05. B) Thresholded maps of voxels with significant coupling associations with sex after FDR correction at 0.05.

### 3.3 CBF-ALFF-ReHo coupling varies between males and females, primarily in subcortical regions

Associations between CBF-ALFF-ReHo coupling and sex were present primarily in the hippocampus and thalamus (Figure 3; corrected p < 0.05). Sex differences in coupling were rare in other subcortical structures and all functional networks – only 1% to 3% of voxels in these regions showed coupling and sex associations (Figure 5). Spin testing between functional networks and sex-related changes in coupling revealed enrichment of coupling and sex associations in the frontoparietal network (p = 0.012), despite the small proportion of the frontoparietal network that exhibited coupling associations with sex.

### 3.4 pIMCo provides a consistent estimator of local coupling compared to WLR-based IMCo

In 2016, Vandekar et al. introduced a method to study IMCo relationships at the single voxel level based on local weighted linear regression (WLR) slopes (Vandekar et al., 2016). Because this method relies on estimating WLR slopes between modalities, it is inherently limited to the two-modality setting, cannot account for statistical relationship between modalities, and requires specification of one modality as the independent variable, leading to asymmetry. These limitations are demonstrated by a two-modality example in Figure 6. We see that, if ALFF is defined as the independent variable, the WLR coupling value is 0.25, indicating little coupling, but if CBF is defined as the independent variable, the WLR coupling value is 1.25, indicating five times as much coupling. Thus, when using WLR-based IMCo, two possible coupling values exist at every voxel, and there isno guarantee analyses will show comparable findings between the two. Additionally, these WLR coupling values only describe the trend of the relationship between ALFF and CBF, but do not account for the statistical strength of that relationship.

**Figure 6:**
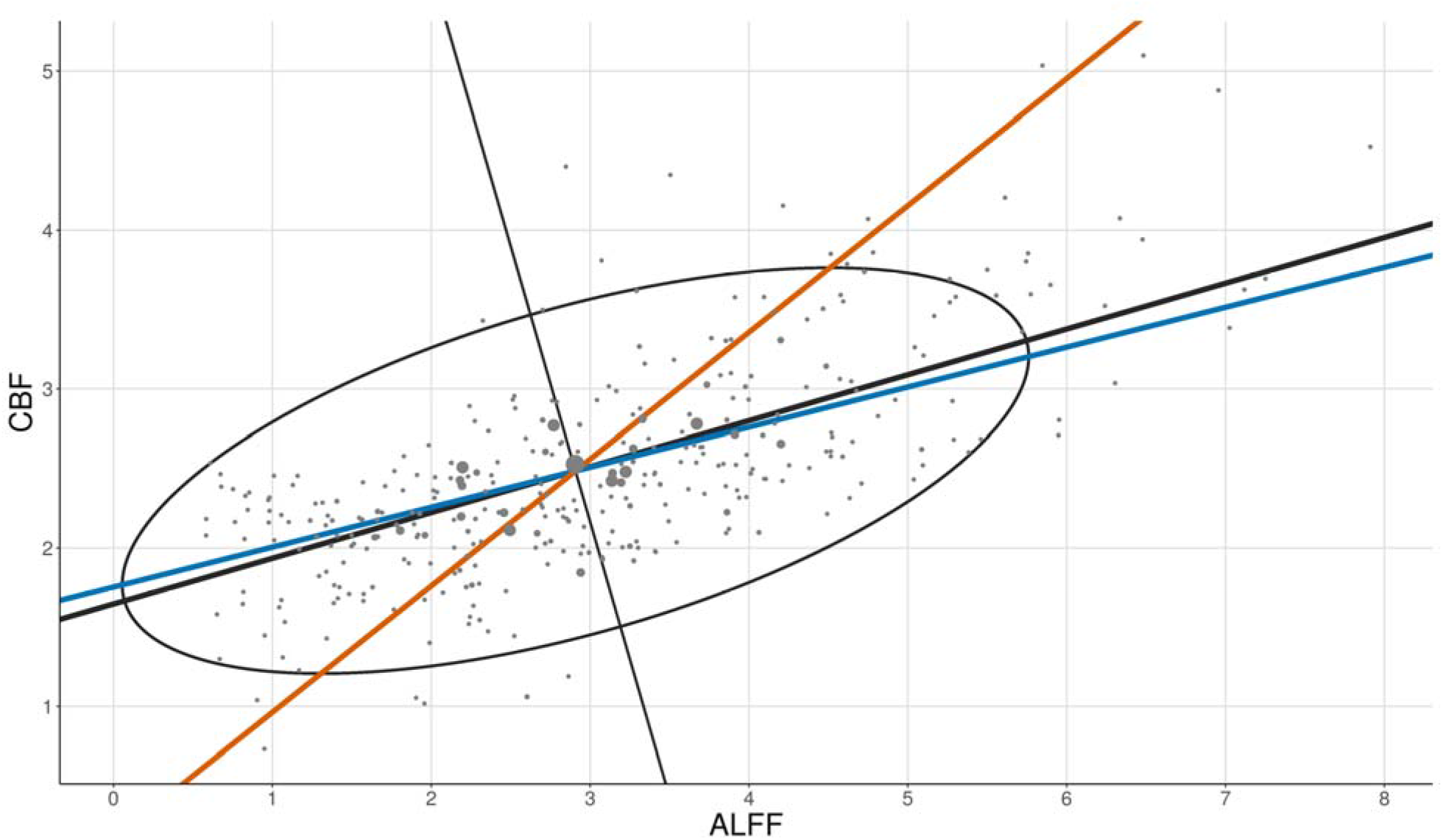
Two-modality example showing pIMCo results in comparison to WLR-based IMCo results. Blue line represents coupling value from WLR-based IMCo if ALFF is defined as the independent variable (slope = 0.25). Orange line represents coupling value from WLR-based IMCo if CBF is defined as the independent variable (slope = 1.25). Black line and ellipse represent PCA results; no reference specification is needed (coupling value = 2.13). Larger dot sizes correspond to increased weights in the WLR and weighted PCA.

In contrast, the pIMCo coupling value is 2.13 and does not require specification of which modality is treated as the independent variable, leading to a symmetric and consistent definition of coupling. This pIMCo value does not describe the effect size of the relationship between ALFF and CBF and instead describes the strength and shape of the relationship – a high value of coupling suggests that the shape of data from that neighborhood looks like a long ellipsoid, while a low value of coupling suggests that the shape is more spherical.

## 4 Discussion

As growing emphasis is placed on the acquisition of multi-modal data, new methodologies are necessary to enable these analyses. In this manuscript, we introduce pIMCo, a generalized approach to estimating local IMCo that can be applied to two or more modalities, can be interpreted as a direct summary of local covariance matrices, and is symmetric (Figure 6). Our method can be used with any combination of volumetric images to produce single-subject, voxel-resolution coupling images which can then be analyzed using standard techniques. We applied our proposed method to show significant coupling between cerebral blood flow, resting state fluctuations, and local connectivity throughout the brain. We then used voxel-level analyses to characterize how coupling varies in neurodevelopment.

### 4.1 Coupling of cerebral blood flow, resting state fluctuations, and local connectivity vary with age and sex

We found that CBF-ALFF-ReHo coupling was spatially heterogeneous and varied with age and sex in neurodevelopment in both subcortical structures and functional networks.These findings, which uncover otherwise undetectable intermodal interactions, are unique to those from individual modality analyses.

We noticed that regions with higher CBF-ALFF-ReHo coupling across subjects also tended to have higher variance in coupling. This suggests that these regions may be biologically interesting in the context of CBF-ALFF-ReHo coupling and important regions for future exploration, since they appear to demonstrate pronounced differences in coupling phenotypes between subjects and could be associated with other variables of interest, such as clinical phenotypes.

Spin testing showed that the high proportion of coupling-age associations in the frontoparietal and default networks were enriched when compared to the cortex overall. This suggests that in neurodevelopment, there is change in not only blood flow, resting state fluctuations, and local connectivity individually, but also in the strength of interaction among these features. These findings are consistent with and fortify the literature demonstrating the importance of frontoparietal and default networks as regions for change in neurodevelopment (Baller et al., 2021; Chai et al., 2017; Fair et al., 2008; Lin et al., 2019). Outside of neurodevelopment, our findings are consistent with previous work showing that coupling between CBF and functional connectivity strength is stronger in frontoparietal and default networks than regions outside these networks (Tak et al., 2015). In subcortical structures, high proportions of coupling associations with age seen in the caudate, pallidum, hippocampus, and thalamus suggest that modulation of vascular, resting state fluctuations, and local connectivity coupling may be necessary in the development of movement, memory, and fundamental brain activities.

High proportions of coupling associations with sex in the hippocampus and thalamus suggest that male-female differences in memory and related cognitive functions between males and females could be explained in part by the strength of regional brain metabolism as measured by cerebral blood flow, resting state fluctuations, and local connectivity coupling. In the cortex, the rarity of coupling-sex associations suggests that this coupling may not play a role in explaining cortical sex-based differences or that this relationship is more complex than our analyses could uncover. These cortical findings are of interest when compared to previous work showing a high proportion of the cortex had significant associations between sex and CBF-ALFF coupling (Baller et al., 2021). Together, these studies demonstrate that three-modality coupling identifies unique relationships when compared to two-modality coupling.

Notably, despite low coupling-sex signal in the frontoparietal network, spin testing showed enrichment of coupling associations with sex in this network. Since spin testing is a spatial permutation test that uses the coupling-sex association thresholded p-value map to generate data under the null, this significant finding is likely due to a combination of overall rare coupling-sex associations in the cortex and the particular spatial distribution of these associations within the frontoparietal network. This finding highlights potential shortcomings of using the spin test for enrichment analysis. Additionally, statistical analysis of enrichment in subcortical structures is not yet possible, so more methods development is needed in this area.

### 4.2 Limitations and Future Directions

pIMCo is designed to summarize a complex local covariance structure, which necessarily leads to outputs that cannot fully characterize the intricacies of coupling. This is especially true in settings with more than two modalities – the covariance structure becomes even more complex and more information is lost when it is summarized. As such, high coupling values can result from many different covariance features, and it is challenging to understand the basis of these high values.

The symmetric nature of pIMCo is conducive to more consistent interpretation when compared to slope-based IMCo, since pIMCo does not depend on reference modality specification. However, while slope is an biologically intuitive measure that can be interpreted as capturing the directionality and effect size of a relationship, the generalizable nature of the pIMCo value makes it challenging to interpret biologically, especially when used on more than two modalities. Instead, it is most accurately interpreted in statistical terms – as a measure of the proportion of variance explained by the first eigenvector, as a measure of how well the local covariance structure could be summarized in one dimension, or as a measure of how ellipsoidal the neighborhood is instead of spherical. This is shown in Figure 6, where the WLR coupling value estimates that, on average, a 1 unit increase in CBF corresponds to a 1.25 unit increase in ALFF, while the pIMCo coupling value describes the ratio of the ellipse’s major axis to minor axis and suggests the statistical strength of the relationship between ALFF and CBF is strong.

Next, since PCA functions as a linear dimension reduction technique, it is most effective at summarizing data whose shape is roughly ellipsoid. However, it is unlikely that such an assumption holds for all voxels and all combinations of different modalities. For example, in a two-dimensional neighborhood, it could be possible that there is a strong quadratic relationship between modalities, but the data is not well-summarized by one eigenvalue and its corresponding eigenvector. In such cases, an IMCo technique based on manifold learning concepts could be a useful improvement that picks up on otherwise undetected intermodal relationships.

Finally, pIMCo is designed to estimate coupling in cross-sectional multi-modality datasets. However, there is also rich covariance information in longitudinal datasets with one or multiple modalities. Future implementations of pIMCo suitable for longitudinal data could investigate whether IMCo changes over time in neurodevelopment or disease, and if so, whether such changes may be useful in identifying biomarkers for clinical phenotypes.

## 5 Conclusion

pIMCo offers a novel perspective for summarizing the overall covariance structure between more than two modalities as well as a generalized, symmetric approach for describing coupling in the two-modality setting. Here, we applied this method to the analysis of coupling between cerebral blood flow, resting state fluctuations, and local connectivity images. This analysis revealed patterns in neurodevelopment with respect to age and sex that differed from those present in any individual modality. As multi-modal data becomes more common, we hope that pIMCo will serve as a tool for capturing complex intermodal relationships and enable more sophisticated descriptive analyses, improved prediction efforts, and novel methodological advances, among others.

## Acknowledgements

This study was supported by grants from the National Institute of Mental Health (R01MH112847 to TDS and RTS; R01MH120482 and R01MH113550 to TDS; R01 MH123550-01 to RTS; R01MH107235 to RCG; 2T32MH019112-29A1 to EBB; R01MH120174, R01MH119185, and R56AG066656 to DRR). FH was supported by NIH Medical Scientist Training Program T32 GM07170. SMW was supported by the National Science Foundation Graduate Research Fellowship Program (NSF GRFP). The PNC data used was funded by NIMH RC2 grants (MH089983 and MH089924 to REG). Funding sources were not involved in study design, data analysis, manuscript preparation, or submission decisions.

## 6 Supplementary Materials

**Figure S1:**
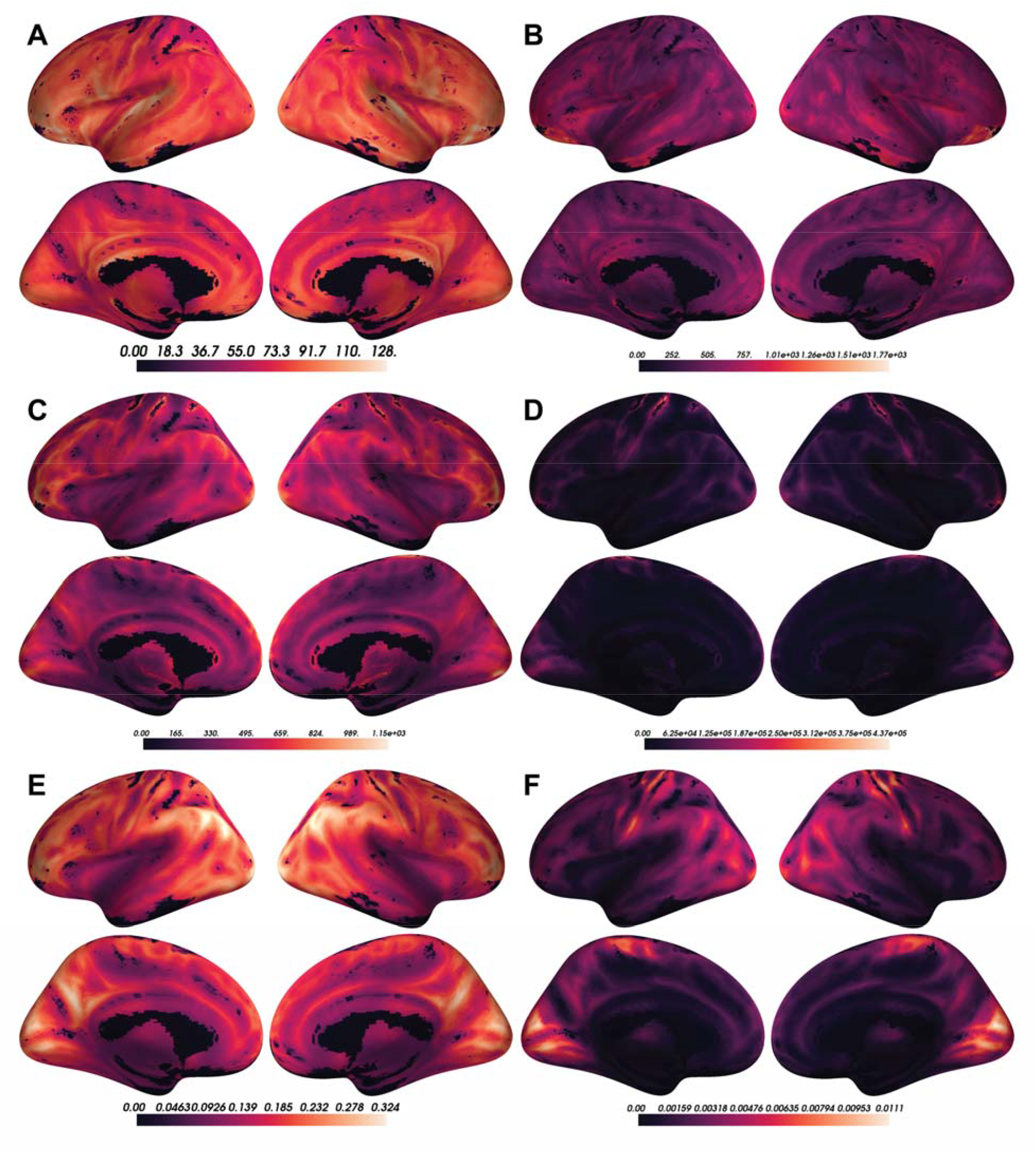
A) Voxel-wise means across subjects of cortical CBF. B) Voxel-wise variances across subjects of cortical CBF. C) Voxel-wise means across subjects of cortical ALFF. D) Voxel-wise variances across subjects of cortical ALFF. E) Voxel-wise means across subjects of cortical ReHo. F) Voxel-wise variances across subjects of cortical ReHo.

## References

Alexander-Bloch, A.F., Shou, H., Liu, S., Satterthwaite, T.D., Glahn, D.C., Shinohara, R.T., Vandekar, S.N., Raznahan, A., 2018. On testing for spatial correspondence between maps of human brain structure and function. NeuroImage 178, 540–551. https://doi.org/10.1016/j.neuroimage.2018.05.070

Allaire, J., Xie, Y., McPherson, J., Luraschi, J., Ushey, K., Atkins, A., Wickham, H., Cheng, J., Chang, W., Iannone, R., 2021. Rmarkdown: Dynamic documents for r.

Avants, B.B., 2020. ANTsR: ANTs in r: Quantification tools for biomedical images.

Avants, B.B., Tustison, N.J., Song, G., Cook, P.A., Klein, A., Gee, J.C., 2011. A reproducible evaluation of ANTs similarity metric performance in brain image registration. NeuroImage 54, 2033–2044. https://doi.org/10.1016/j.neuroimage.2010.09.025

Baller, E.B., Valcarcel, A.M., Adebimpe, A., Alexander-Bloch, A., Cui, Z., Gur, R.C., Gur, R.E., Larsen, B.L., Linn, K.A., O’NADonnell, C.M., Pines, A.R., Raznahan, A., Roalf, D.R., Sydnor, V.J., Tapera, T.M., Tisdall, M.D., Vandekar, S., Xia, C.H., Detre, J.A., Shinohara, R.T., Satterthwait, T.D., 2021. Developmental coupling of cerebral blood flow and fMRI fluctuations in youth. bioRxiv. https://doi.org/10.1101/2021.07.28.454179

Benjamini, Y., Hochberg, Y., 1995. Controlling the False Discovery Rate: A Practical and Powerful Approach to Multiple Testing. Journal of the Royal Statistical Society. Series B (Methodological) 57, 289–300.

Chai, L.R., Khambhati, A.N., Ciric, R., Moore, T.M., Gur, R.C., Gur, R.E., Satterthwaite, T.D., Bassett, D.S., 2017. Evolution of brain network dynamics in neurodevelopment. Network Neuroscience 1, 14–30. https://doi.org/10.1162/NETN_a_00001

Chappell, M.A., Groves, A.R., MacIntosh, B.J., Donahue, M.J., Jezzard, P., Woolrich, M.W., 2011. Partial volume correction of multiple inversion time arterial spin labeling MRI data. Magnetic Resonance in Medicine 65, 1173–1183. https://doi.org/10.1002/mrm.22641

Chappell, M.A., Groves, A.R., Whitcher, B., Woolrich, M.W., 2009. Variational Bayesian Inference for a Nonlinear Forward Model. IEEE Transactions on Signal Processing 57, 223–236. https://doi.org/10.1109/TSP.2008.2005752

Ciric, R., Rosen, A.F., Erus, G., Cook, P.A., Bassett, D.S., Davatzikos, C., Wolf, D.H., Satterthwaite, T.D., 2018. Mitigating head motion artefact in functional connectivity MRI. Nat Protoc 13, 2801–2826. https://doi.org/10.1038/s41596-018-0065-y

Ciric, R., Thompson, W.H., Lorenz, R., Goncalves, M., MacNicol, E., Markiewicz, C.J., Halchenko, Y.O., Ghosh, S.S., Gorgolewski, K.J., Poldrack, R.A., Esteban, O., 2021. TemplateFlow: FAIR-sharing of multi-scale, multi-species brain models 2021.02.10.430678. https://doi.org/10.1101/2021.02.10.430678

Ciric, R., Wolf, D.H., Power, J.D., Roalf, D.R., Baum, G.L., Ruparel, K., Shinohara, R.T., Elliott, M.A., Eickhoff, S.B., Davatzikos, C., Gur, R.C., Gur, R.E., Bassett, D.S., Satterthwaite, T.D., 2017. Benchmarking of participant-level confound regression strategies for the control of motion artifact in studies of functional connectivity. NeuroImage, Cleaning up the fMRI time series: Mitigating noise with advanced acquisition and correction strategies 154, 174–187. https://doi.org/10.1016/j.neuroimage.2017.03.020

Cox, R.W., 1996. AFNI: Software for analysis and visualization of functional magnetic resonance neuroimages. Computers and Biomedical research 29, 162–173.

Desikan, R.S., S’egonne, F., Fischl, B., Quinn, B.T., Dickerson, B.C., Blacker, D., Buckner, R.L., Dale, A.M., Maguire, R.P., Hyman, B.T., Albert, M.S., Killiany, R.J., 2006. An automated labeling system for subdividing the human cerebral cortex on MRI scans into gyral based regions of interest. Neuroimage 31, 968–980. https://doi.org/10.1016/j.neuroimage.2006.01.021

Fair, D.A., Cohen, A.L., Dosenbach, N.U.F., Church, J.A., Miezin, F.M., Barch, D.M., Raichle, M.E., Petersen, S.E., Schlaggar, B.L., 2008. The maturing architecture of the brain’s default network. PNAS 105, 4028–4032. https://doi.org/10.1073/pnas.0800376105

Fischl, B., 2012. FreeSurfer. Neuroimage 62, 774–781.

Greve, D.N., Fischl, B., 2009. Accurate and robust brain image alignment using boundary-based registration. NeuroImage 48, 63–72. https://doi.org/10.1016/j.neuroimage.2009.06.060

Gu, Z., Jamison, K.W., Sabuncu, M.R., Kuceyeski, A., 2021. Heritability and interindividual variability of regional structure-function coupling. Nat Commun 12, 4894. https://doi.org/10.1038/s41467-021-25184-4

Gur, R.C., Butler, E.R., Moore, T.M., Rosen, A.F.G., Ruparel, K., Satterthwaite, T.D., Roalf, D.R., Gennatas, E.D., Bilker, W.B., Shinohara, R.T., Port, A., Elliott, M.A., Verma, R., Davatzikos, C., Wolf, D.H., Detre, J.A., Gur, R.E., 2020. Structural and Functional Brain Parameters Related to Cognitive Performance Across Development: Replication and Extension of the Parieto-Frontal Integration Theory in a Single Sample. Cereb Cortex 31, 1444–1463. https://doi.org/10.1093/cercor/bhaa282

Honey, C.J., Sporns, O., Cammoun, L., Gigandet, X., Thiran, J.P., Meuli, R., Hagmann, P., 2009. Predicting human resting-state functional connectivity from structural connectivity. PNAS 106, 2035–2040. https://doi.org/10.1073/pnas.0811168106

Jenkinson, M., Beckmann, C.F., Behrens, T.E., Woolrich, M.W., Smith, S.M., 2012. FSL. Neuroimage 62, 782–790.

Kandel, B.M., Cook, P.A., Tustison, N.J., Muschelli, J., 2020. ANTsRCore: Core software infrastructure for ‘ANTsR’.

Lin, H.-Y., Perry, A., Cocchi, L., Roberts, J.A., Tseng, W.-Y.I., Breakspear, M., Gau, S.S.-F., 2019. Development of frontoparietal connectivity predicts longitudinal symptom changes in young people with autism spectrum disorder. Transl Psychiatry 9, 1–10. https://doi.org/10.1038/s41398-019-0418-5

Müller, K., 2020. Here: A simpler way to find your files.

Muschelli, J., 2021a. Extrantsr: Extra functions to build on the ‘ANTsR’ package.

Muschelli, J., 2021b. Neurobase: ‘Neuroconductor’ base package with helper functions for ‘nifti’ objects.

Muschelli, J., Sweeney, E., Lindquist, M., Crainiceanu, C., 2015. Fslr: Connecting the FSL software with r. The R Journal 7, 163–175.

R Core Team, 2021. R: A language and environment for statistical computing. R Foundation for Statistical Computing, Vienna, Austria.

Ren, K., 2021. Rlist: A toolbox for non-tabular data manipulation.

Satterthwaite, T.D., Elliott, M.A., Ruparel, K., Loughead, J., Prabhakaran, K., Calkins, M.E., Hopson, R., Jackson, C., Keefe, J., Riley, M., Mensh, F.D., Sleiman, P., Verma, R., Davatzikos, C., Hakonarson, H., Gur, R.C., Gur, R.E., 2014. Neuroimaging of the Philadelphia Neurodevelopmental Cohort. NeuroImage 86, 544–553. https://doi.org/10.1016/j.neuroimage.2013.07.064

Schäfer, T., 2021. Freesurferformats: Read and write ‘FreeSurfer’ neuroimaging file formats.

Shokri-Kojori, E., Tomasi, D., Alipanahi, B., Wiers, C.E., Wang, G.-J., Volkow, N.D., 2019. Correspondence between cerebral glucose metabolism and BOLD reveals relative power and cost in human brain. Nat Commun 10, 690. https://doi.org/10.1038/s41467-019-08546-x

Tak, S., Polimeni, J.R., Wang, D.J.J., Yan, L., Chen, J.J., 2015. Associations of Resting-State fMRI Functional Connectivity with Flow-BOLD Coupling and Regional Vasculature. Brain Connect 5, 137–146. https://doi.org/10.1089/brain.2014.0299

Thomas Yeo, B.T., Krienen, F.M., Sepulcre, J., Sabuncu, M.R., Lashkari, D., Hollinshead, M., Roffman, J.L., Smoller, J.W., Zöllei, L., Polimeni, J.R., Fischl, B., Liu, H., Buckner, R.L., 2011. The organization of the human cerebral cortex estimated by intrinsic functional connectivity. J Neurophysiol 106, 1125–1165. https://doi.org/10.1152/jn.00338.2011

Tustison, N.J., Cook, P.A., Klein, A., Song, G., Das, S.R., Duda, J.T., Kandel, B.M., van Strien, N., Stone, J.R., Gee, J.C., Avants, B.B., 2014. Large-scale evaluation of ANTs and FreeSurfer cortical thickness measurements. NeuroImage 99, 166–179. https://doi.org/10.1016/j.neuroimage.2014.05.044

Tzourio-Mazoyer, N., Landeau, B., Papathanassiou, D., Crivello, F., Etard, O., Delcroix, N., Mazoyer, B., Joliot, M., 2002. Automated anatomical labeling of activations in SPM using a macroscopic anatomical parcellation of the MNI MRI single-subject brain. Neuroimage 15, 273–289. https://doi.org/10.1006/nimg.2001.0978

Uddin, L.Q., 2013. Complex relationships between structural and functional brain connectivity. Trends in Cognitive Sciences, Special Issue: The Connectome 17, 600– 602. https://doi.org/10.1016/j.tics.2013.09.011

Valcarcel, A.M., Linn, K.A., Khalid, F., Vandekar, S.N., Tauhid, S., Satterthwaite, T.D., Muschelli, J., Martin, M.L., Bakshi, R., Shinohara, R.T., 2018a. A dual modeling approach to automatic segmentation of cerebral T2 hyperintensities and T1 black holes in multiple sclerosis. NeuroImage: Clinical 20, 1211–1221. https://doi.org/10.1016/j.nicl.2018.10.013

Valcarcel, A.M., Linn, K.A., Vandekar, S.N., Satterthwaite, T.D., Muschelli, J., Calabresi, P.A., Pham, D.L., Martin, M.L., Shinohara, R.T., 2018b. MIMoSA: An Automated Method for Intermodal Segmentation Analysis of Multiple Sclerosis Brain Lesions. J Neuroimaging 28, 389–398. https://doi.org/10.1111/jon.12506

Vandekar, S.N., Shinohara, R.T., Raznahan, A., Hopson, R.D., Roalf, D.R., Ruparel, K., Gur, R.C., Gur, R.E., Satterthwaite, T.D., 2016. Subject-level Measurement of Local Cortical Coupling. Neuroimage 133, 88–97. https://doi.org/10.1016/j.neuroimage.2016.03.002

Waskom, M., Gramfort, A., Burns, S., Luessi, M., Larson, E., 2020. PySurfer.

Wickham, H., 2019. Stringr: Simple, consistent wrappers for common string operations.

Wickham, H., Averick, M., Bryan, J., Chang, W., McGowan, L.D., François, R., Grolemund, G., Hayes, A., Henry, L., Hester, J., Kuhn, M., Pedersen, T.L., Miller, E., Bache, S.M., Müller, K., Ooms, J., Robinson, D., Seidel, D.P., Spinu, V., Takahashi, K., Vaughan, D., Wilke, C., Woo, K., Yutani, H., 2019. Welcome to the tidyverse. Journal of Open Source Software 4, 1686. https://doi.org/10.21105/joss.01686

Xie, Y., 2021a. Bookdown: Authoring books and technical documents with r markdown.

Xie, Y., 2021b. Knitr: A general-purpose package for dynamic report generation in r.

Xie, Y., 2016. Bookdown: Authoring books and technical documents with R markdown. Chapman; Hall/CRC, Boca Raton, Florida.

Yang, H., Long, X.-Y., Yang, Y., Yan, H., Zhu, C.-Z., Zhou, X.-P., Zang, Y.-F., Gong, Q.-Y., 2007. Amplitude of low frequency fluctuation within visual areas revealed by resting-state functional MRI. Neuroimage 36, 144–152. https://doi.org/10.1016/j.neuroimage.2007.01.054

Yushkevich, P.A., Piven, J., Cody Hazlett, H., Gimpel Smith, R., Ho, S., Gee, J.C., Gerig, G., 2006. User-guided 3D active contour segmentation of anatomical structures: Significantly improved efficiency and reliability. Neuroimage 31, 1116–1128.

Zang, Y., Jiang, T., Lu, Y., He, Y., Tian, L., 2004. Regional homogeneity approach to fMRI data analysis. Neuroimage 22, 394–400. https://doi.org/10.1016/j.neuroimage.2003.12.030

